# Functionally Important Residues from Graph Analysis of Coevolved Dynamic couplings

**DOI:** 10.1101/2024.10.30.621065

**Authors:** Manming Xu, Sarath Chandra Dantu, James A Garnett, Robert A Bonomo, Alessandro Pandini, Shozeb Haider

**Author notes:** Joint first authors (Manming Xu, Sarath Chandra Dantu). To whom correspondence should be addressed: Prof Shozeb Haider, Dr Alessandro Pandini.

## Abstract

The relationship between protein dynamics and function is essential for understanding biological processes and developing effective therapeutics. Functional sites within proteins are critical for activities such as substrate binding, catalysis, and structural changes. Existing computational methods for the predictions of functional residues are trained on sequence, structural and experimental data, but they do not explicitly model the influence of evolution on protein dynamics. This overlooked contribution is essential as it is known that evolution can fine tune protein dynamics through compensatory mutations, either to improve the proteins’ performance or diversify its function while maintaining the same structural scaffold. To model this critical contribution, we introduce DyNoPy, a computational method that combines residue coevolution analysis with molecular dynamics (MD) simulations, revealing hidden correlations between functional sites. DyNoPy constructs a graph model of residue-residue interactions, identifies communities of key residue groups and annotates critical sites based on their roles. By leveraging the concept of coevolved dynamical couplings—residue pairs with critical dynamical interactions that have been preserved during evolution—DyNoPy offers a powerful method for predicting and analysing protein evolution and dynamics. We demonstrate the effectiveness of DyNoPy on SHV-1 and PDC-3, chromosomally encoded β-lactamases linked to antibiotic resistance, highlighting its potential to inform drug design and address pressing healthcare challenges.

## Introduction

Quantifying the contribution of individual residues or residue groups to protein function is important to estimate the pathogenic effect of mutations (1). Identifying the functional roles of individual residues has primarily been done through mutagenesis experiments (2). Bioinformatics methods have complemented these approaches through analysis of multiple sequence alignments (MSA) of homologous proteins and structural data (3–8). Among these methods, computational techniques that can decode inter-residue evolutionary relationships from MSAs have paved the way for machine learning (ML) based strategies that can predict protein structure (9–12), stability (13), and function (7) and extend the scope of computational protein design (14–16). A most recent approach has combined experimental data from three proteins, NUDT15, PTEN and CYP2C9, on stability and function with sequence and structural features to train a ML model to predict functional sites (17).

Functional sites are often regulated by both, local and global interactions. Changes in these interactions are instrumental for functional events like substrate binding, catalysis, and conformational changes (18). The development of physical models of protein dynamics and the increase in available computational power has stimulated the adoption of computational techniques (19, 20) to investigate the conformational dynamics of proteins, an essential component of the many biological functions (21, 22). Different models have been proposed to describe the interactions between residues during simulations and network models have been particularly popular, including methods on single structures and MD simulations data built by analysing the response to external forces on residue networks (23), by estimating the prevalence of non-covalent energy interaction networks in homologous proteins (24), or by analysing linear or non-linear correlation in atomic fluctuations (25, 26). These techniques have demonstrated their usefulness in extracting allosteric networks from structural data with applications in enzyme design (26).

However, none of these techniques incorporate information on residue evolution into the computational approach, While it has been established that evolution through compensatory mutations in dynamic regions, like hinges and loops, can fine tune protein structural dynamics and introduce promiscuity, thereby diversifying biological function. Assuming that protein functional dynamics is conserved during evolution, significant information on dynamic regions and substrate recognition sites should be recoverable using inter residue coevolution scores extracted from MSAs (27, 28). Coevolution analysis and Molecular Dynamics (MD) simulations have independently (29) and synergistically been combined in the past to identify important residues for function (30–34). Yet a method that combines hidden information on dynamics from evolution with direct information on local and global dynamics from conformational ensembles from MD is not yet available.

Here, we present DyNoPy, a computational method that can extract hidden information on functional sites from the combination of pairwise residue coevolution data and powerful descriptors of dynamics extracted from the analysis of MD ensembles. The method can detect coevolved dynamic couplings, i.e. residue pairs with critical dynamical interactions that have been preserved during evolution. These pairs are extracted from a graph model of residue-residue interactions. Communities of important residue groups are detected, and critical sites are identified by their eigenvector centrality in the graph (Fig. 1). We demonstrate the power of this approach on SHV-1 and PDC-3 β-lactamases of major clinical importance (35, 36). DyNoPy successfully detects residue couplings that align with previous studies, guide in the explanations of mutation sites with previously unexplained mechanisms and provide predictions on plausible important sites for the emergence of clinically relevant variants.

**Fig. 1 –.**
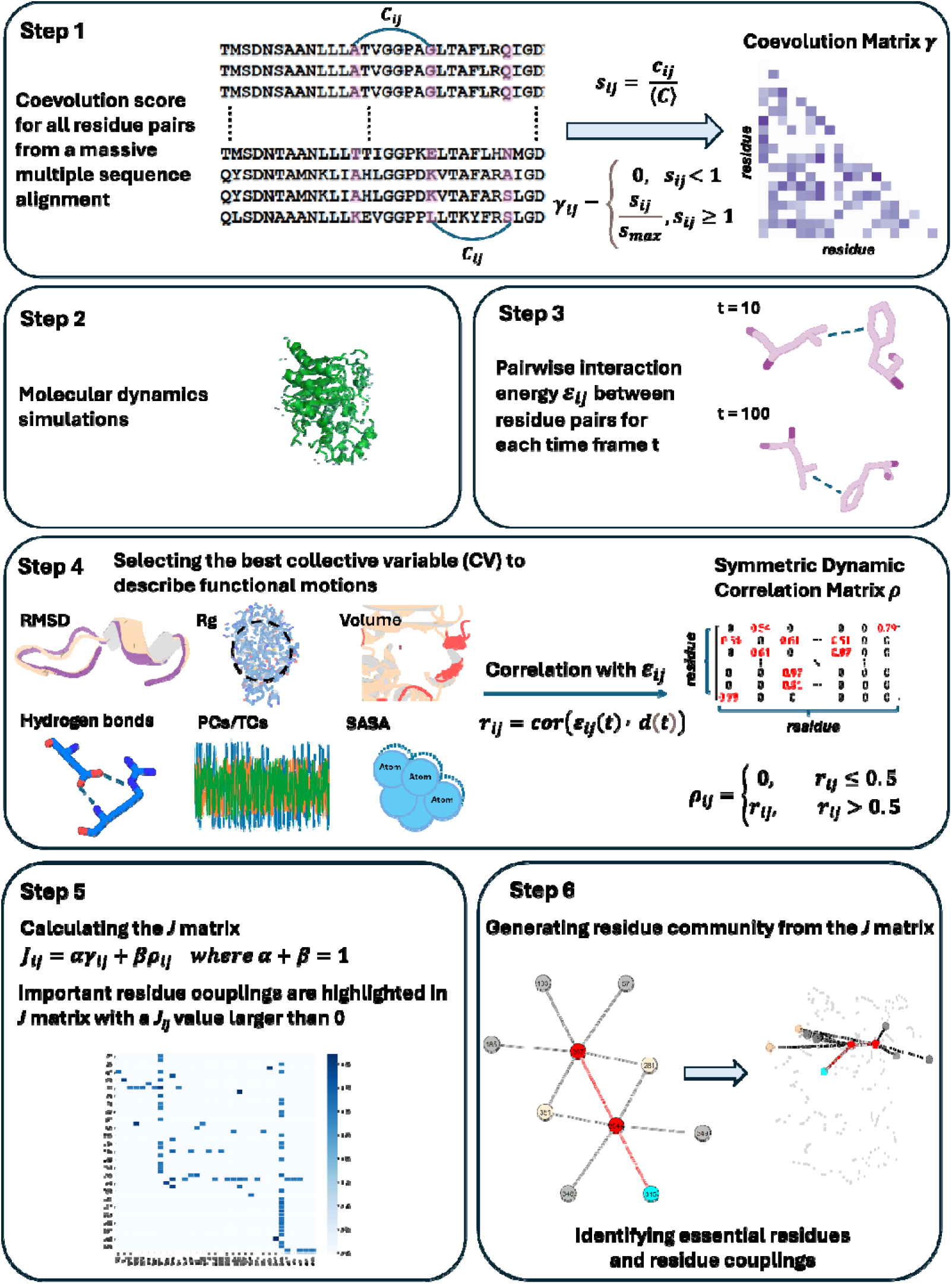
Overview of the DyNoPy workflow

## Results and Discussion

β-lactamases are a group of enzymes capable of hydrolysing β-lactams, conferring resistance to β-lactam antibiotics (37). These enzymes are evolving rapidly, as single amino acid substitutions are sufficient to drive their evolution and increase their catalytic spectrum and inhibitor resistance profile (38). The widespread dissemination of β-lactamases across different bacterial species and their extensive emergence highlight their global impact on antibiotic resistance (39). The rapid evolution of β-lactamases and their clinical significance (38) makes them an ideal target for evaluating the robustness of DyNoPy.

In this study, we applied DyNoPy to two model enzymes from different β-lactamase families: class A β-lactamase SHV-1 (a chromosomally encoded enzyme in *Klebsiella pneumoniae*) and class C β-lactamase PDC-3 (a chromosomally encoded enzyme in *Pseudomonas aeruginosa*) (35, 36) (Supplementary Fig. S1 and S2). Both class A and class C β-lactamases comprise an α/β domain and an α helical domain, with the active site situated in between (40, 41). Moreover, both enzymes target the carbonyl carbon of the β-lactams using a highly conserved serine residue (42, 43). Despite these similarities, the structures of class A and class C β-lactamases are remarkably different (Fig. 2).

**Fig. 2 –.**
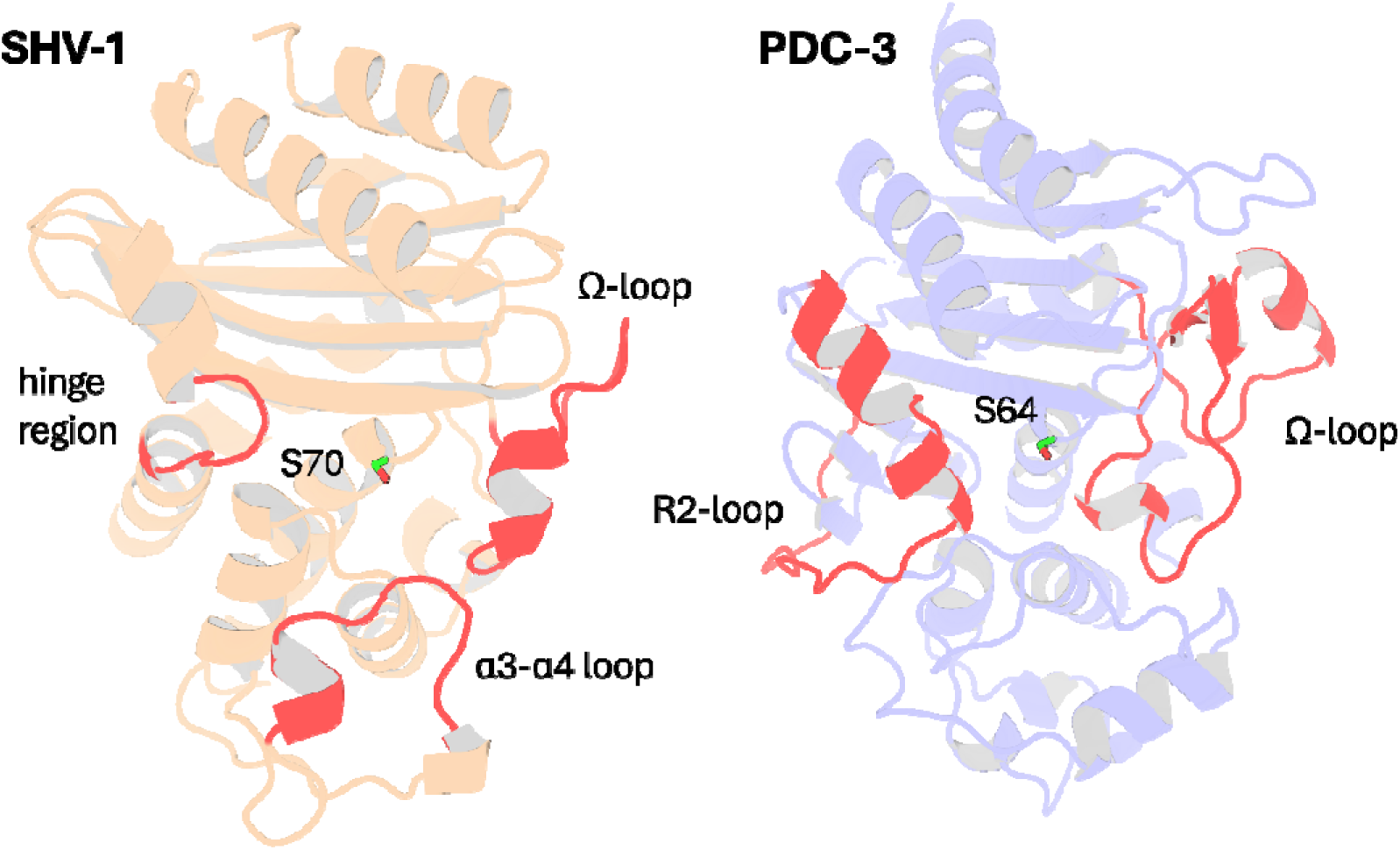
Structural Comparison of SHV-1 (PDB ID: 3N4I) and PDC-3 (PDB ID: 4HEF) β-Lactamases. Catalytic serine S_70_ (SHV-1) and S_64_ (PDC3) are highlighted using stick representation. Important loops surrounding the active site are highlighted in red. In SHV-1, highlighted loops are the α3-α4 loop (residues 101-111), the Ω-loop (residues 164-179), and the hinge region (residues 213-218). In PDC-3, highlighted loops are the Ω-loop (residues 183-226) and the R2-loop (residues 280-310).

SHV-1 is a very well characterised enzyme with wealth of information on mutations and their corresponding effects on protein function. In contrast, the information available on PDC-3 remains limited. Detailed structural information on these enzymes can be found in the supplementary materials. Essential catalytic residues in SHV-1 are: S_70_, K_73_, S_130_, E_166_, N_170_, K_234_, G_236_, and A_237_ (44) and conserved catalytic residues in PDC-3 include S_64_, K_67_, Y_150_, N_152_, K_315_, T_316_, and G_317_. Highly conserved stretches of 3-9 hydrophobic residues, annotated as hydrophobic nodes, exists in class A β-lactamases and have been proven to be essential for protein stability (45). Residues defined as belonging to hydrophobic nodes within SHV-1 are listed in Supplementary Table S1.

In SHV-1, the predominant extended spectrum β-lactamase (ESBL) substitutions occur at L_35_, G_238_, and E_240_, while R_43_, E_64_, D_104_, A_146_, G_156_, D_179_, R_202_, and R_205_ appear in ESBLs with lower frequency (46). Mutations at M_69_, S_130_, A_187_, T_235_, and R_244_ are known to induce inhibitor resistance in the enzyme (47). In PDC-3, substitutions primarily occur on the Ω-loop, enhancing its flexibility to accommodate the bulky side chains of antibiotics, while deletions are more common in the R2-loop (43). The predominant Ω-loop mutations isolated from clinics are found at positions V_211_, G_214_, E_219_, and Y_221_ (48).

### Emergence of highly conserved dynamic couplings

DyNoPy builds a pairwise model of conserved dynamic couplings detected by combining coevolution scores and information on functional motions into a score *J_ij_* (see Methods and Fig. 1). To this end a dynamics descriptor should be selected. When the descriptor is associated with functional conformational changes, it is expected that functionally relevant couplings will report higher scores. Dynamics descriptors can be selected from commonly used geometrical collective variables (CVs) for the analysis of MD trajectories (see Methods). As expected, the average ***J*** matrix score varies across the different CVs, with some of them showing no signal of dynamic coupling (Supplementary Fig. S4C).

SHV-1 and PDC-3 exhibit distinct dynamics, requiring a different choice of the CV that best captures the functional dynamics. For SHV-1, the global first principal component (PC1) proved to be the most effective feature, identifying 571 residue pairs with a *J_ij_* value greater than 0. Conversely, PDC-3 requires selection of more localized features that can extract the Ω-loop dynamics from the overall protein motion. Among the dynamic descriptors, the partial first time-lagged component (TC1_partial) performed best for PDC-3, detecting 216 residue pairs with a *J_ij_* value greater than 0. Consequently, PC1 and TC1_partial were selected to build the ***J*** matrix for SHV-1 and PDC-3, respectively. The performance of all 12 CVs for each protein was assessed and listed in the Supplementary Table S2.

The importance of dynamical information is evident when coevolution couplings (γ*_ij_*) and conserved dynamic couplings (*J_ij_*) are compared: the number of non-zero couplings decrease from 40% to <2% of total residue pairs in the protein (Supplementary Fig. S4D) when information from the dynamics descriptor is added. Thus, the inclusion of protein dynamics in coevolution studies acts as an effective filter that rules out residue pairs that do not have significant correlations with functional motions. Moreover, when relying only on γ*_ij_*, all the residues in SHV-1 and PDC-3 are included within four identified communities (Supplementary Table S3), suggesting that coevolution scores (γ*_ij_*) alone do not effectively discriminate residues relevant for protein functions. Furthermore, it would be hard to distinguish critical core residues for each community using only γ*_ij_*, as the eigenvector centrality (EVC) values for the residues do not show remarkable differences (Supplementary Fig. S5A and S5B). This means that detailed dynamic investigation of the top residues is needed to determine which pairs should be picked up and further analysed. On the other hand, it is much easier to identify essential residues based on *J* scores calculated, as clear outliers with significantly higher EVC values could be seen for almost all communities (Supplementary Fig. S5C and S5D) (29, 49). In conclusion, the lack of specificity in the statistically based coevolution analysis supports the choice of incorporating a score for the correlation between residue interactions and dynamic behaviours that enables deconvolution of community information.

### DyNoPy reveals critical residues and predicts evolutionary pathways in SHV-1

DyNoPy identified eight meaningful communities, each consisting of at least three strongly coupled residues within SHV-1 (Supplementary Fig. S4A). All crucial catalytic residues and critical substitution sites previously mentioned participating in one of these communities with the exceptions of R_43_, R_202_, and S_130_. Residues previously known to have critical role in function or conferring ESBLs/IRBLs phenotype are either directly coupled to protein dynamics or act as a central hub. The hubs interact with residues with either a role in catalysis or structural stability through their membership of hydrophobic nodes (35). Furthermore, DyNoPy identified key positions (L_162_ and N_136_) within some communities that are known to undergo substitutions, conferring an ESBL phenotype in other class A β-lactamases. These substitutions have not yet emerged in the SHV family, providing insightful predictions about the potential future evolution of the enzyme. Detailed description of communities with secondary importance for protein function (community 3, 8, and 9) is provided in the supplementary information (Supplementary Fig. S6).

### DyNoPy predicts mutation hotspots in SHV-1

DyNoPy detects critical mutation sites (L_162_ and N_136_) that are known to extend the range of substrates in other class A β-lactamases but have not yet emerged as variants in the SHV family. These sites have not been modified in SHV family because of their plausible central role within the communities as they are mediating couplings with key functional residues essential for catalytic activity and structural stability, indicating their critical role in protein function and the potential lower mutation rate. These findings provide insightful predictions about the potential future evolution of the enzyme, as well as plausible explanations for why these mutations have not yet appeared.

L_162,_ positioned at the start of the Ω-loop and adjacent to the crucial catalytic residue E_166,_ is assigned as the core residue for community 1 (Fig. 3A). While it remains conserved in SHV family, variants of L_162_ have been isolated in other class A β-lactamase and are known to expand the enzyme catalytic spectrum. Single amino acid substitution at L_162_ can intensify antibiotic resistance in BEL-1 (50), a class A ESBL clinical variant, exhibiting robust resistance to ticarcillin and ceftazidime (51). BEL-2 diverges from BEL-1 by single amino acid substitution (L_162_F) which alters the kinetic properties of the enzyme significantly and increases its affinity towards expanded-spectrum cephalosporins (52). The relationship between L_162_ and protein catalytic functions can be explained using DyNoPy model, as there are couplings with catalytic important residues M_69_, K_73_, E_166_, and K_234_. Moreover, the BEL case has confirmed that L_162_F mutation significantly destabilizes the overall protein structure, highlighting the crucial role of L_162_ in maintaining protein stability (50). DyNoPy accurately identifies the centrality of L_162_ by reporting its connections with 28 backbone residues, including nine hydrophobic node residues critical for protein stability. Among these, five hydrophobic residues are part of the α2 node: V_75_, L_76_, G_78_, V_80_, and L_81_, highlighting the contribution of L_162_ to the stability of the α2 helix (35).

**Fig. 3 –.**
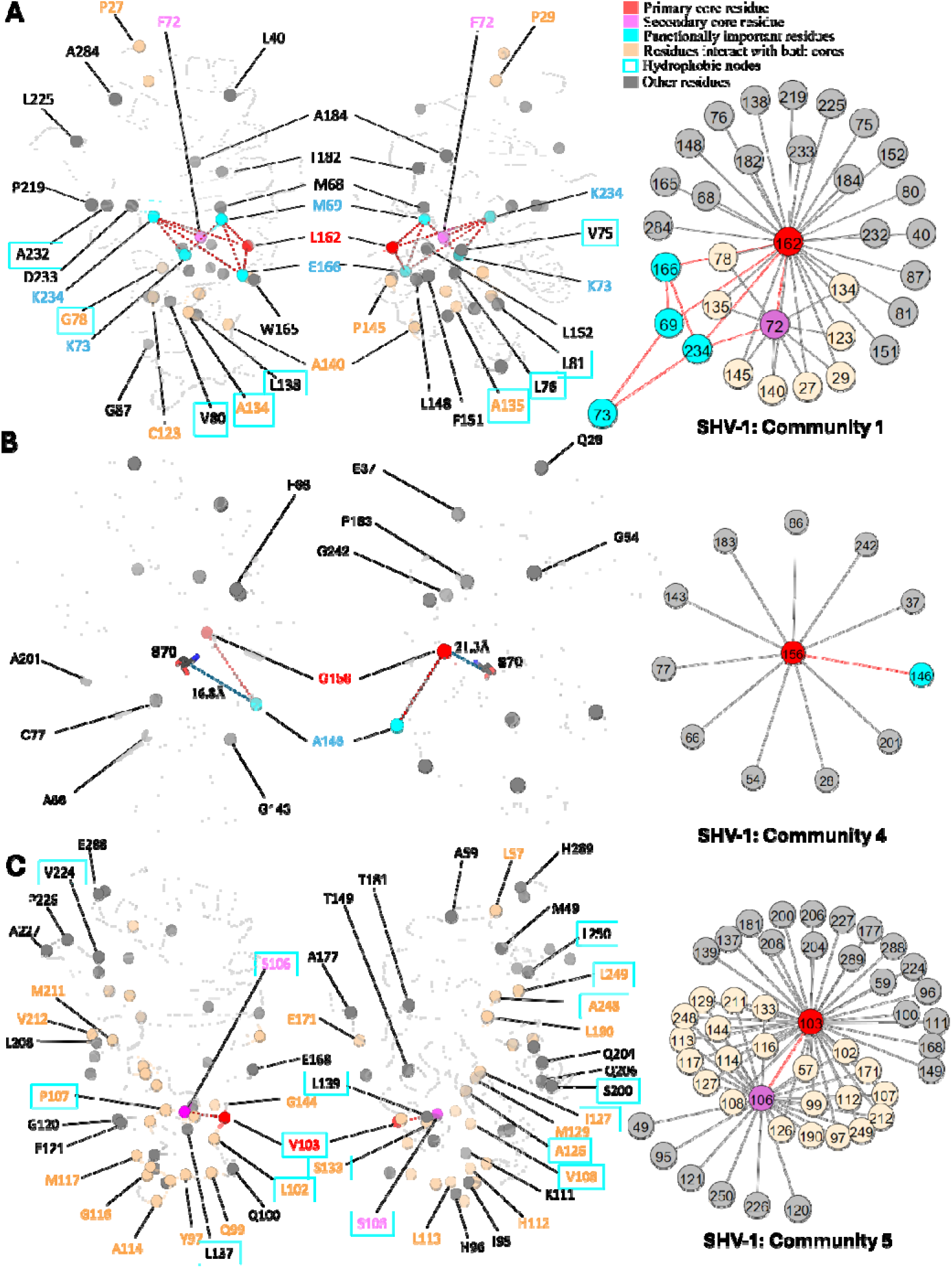
Community 1, 4, and 5 of SHV-1 β-Lactamase. All the residues are depicted as spheres on the protein structure. The core residue for each community is highlighted in red, while purple is used to emphasize the secondary core residue. Residues that interact with both cores are coloured in light yellow. Functional important residues are marked in cyan. Hydrophobic nodes are enclosed with cyan boxes. A. Community 1 of SHV-1, comprising 33 residues with L_162_ being the primary core residue. B. Community 4 of SHV-1, containing 12 residues and is centred by G_156_. G_156_ and A_146_ are two functional important residues distant from the active site. G_156_ is 21.3L away from the catalytic S_70_. A_146_ is 16.8L away from S_70_. C. Community 5 of SHV-1, embracing 48 residues and showing a strong correlation between V_103_ and S_106_.

Just like L_162_, N_136_ undergoes advantageous mutations in other class A β-lactamases while remains highly conserved within the SHV family. It is the core residue for community 7 (Fig. 4B). This residue forms a hydrogen bond with E_166_, stabilizing the Ω-loop (53). Although DyNoPy did not detect this direct interaction between N_136_ and E_166_, the established relationship between N_136_ and N_170_ highlights the role of N_136_ in influencing E_166_. N_170_, an essential catalytic residue located on the Ω-loop, contributes to priming the water molecule for the deacylation step with E_166_ (54) and is directly coupled with N_136_. Due to the essential contribution of N_136_ in facilitating E_166_ to maintain its proper orientation, it was previously thought to be intolerant to mutations as substitution of Asparagine to Alanine at this position would make the enzyme lose its function completely (55). However, N_136_D substitution has emerged as a new clinical variant very recently in PenL, a class A β-lactamase, by increasing its ability in hydrolysing ceftazidime (55), suggesting that this site has potential to mutate. This gain of function is mainly triggered by the increased flexibility of the Ω-loop (55). DyNoPy correctly detect a dynamical relationship between N_136_ and the Ω-loop (residues 164-179). Six residues present in the Ω-loop participate within this community, including R_164_ and D_179_. These two residues are critical as they are forming the ‘bottleneck’ of the Ω-loop which is essential for the correct position of E_166_ (56). D_179_ is also a critical mutation site for SHV-1. Single amino acid substitutions like D_179_A, D_179_N, and D_179_G are enough for the extended spectrum phenotype (46).

**Fig. 4 –.**
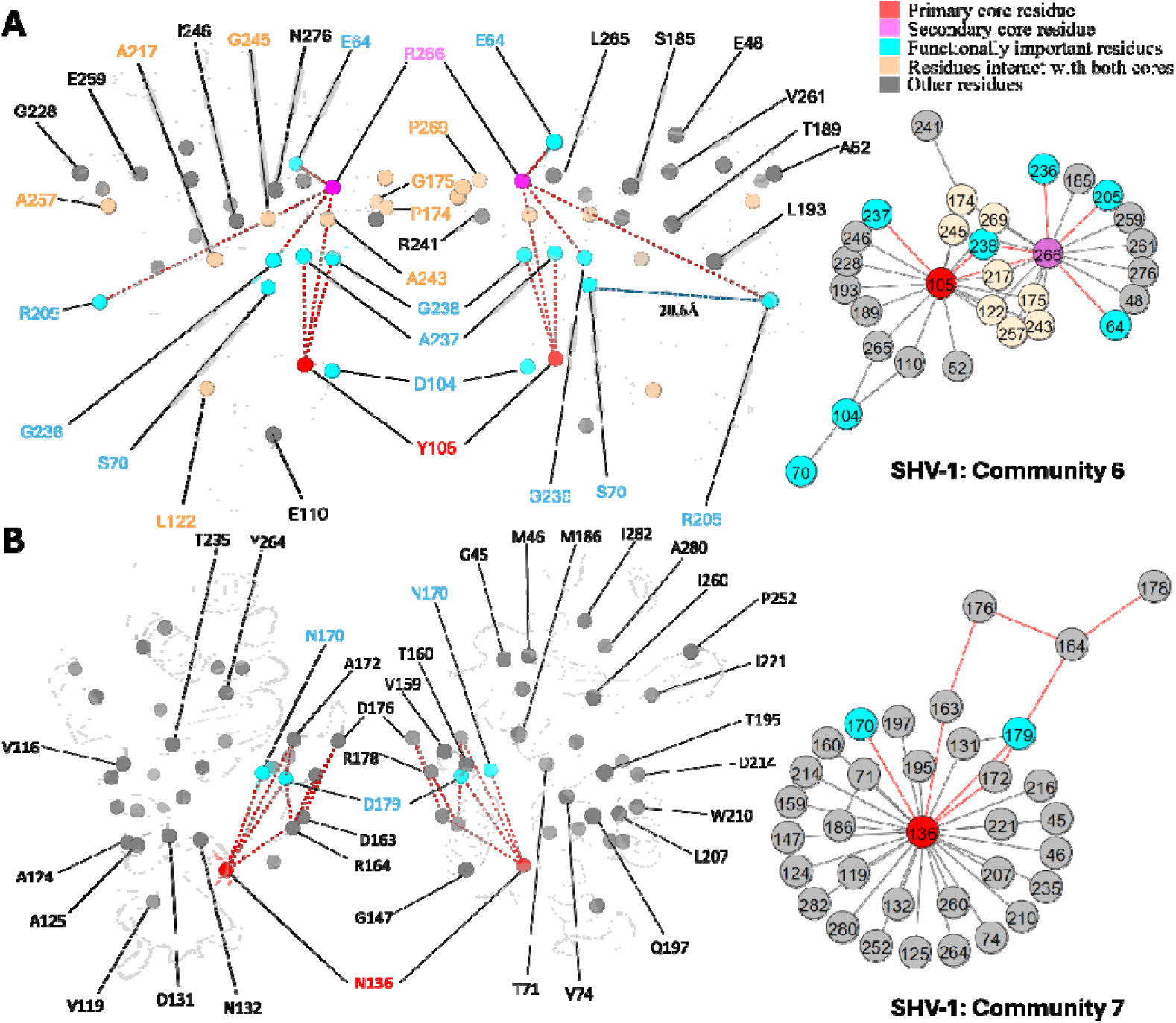
Community 6 and 7 of SHV-1 β-Lactamase. All the residues are depicted as spheres on the protein structure. The core residue for each community is highlighted in red, while purple is used to emphasize the secondary core residue. Residues that interact with both cores are coloured in light yellow. Functional important residues are marked in cyan. A. Community 6 of SHV-1, comprising 30 residues with Y_105_ being the primary core residue. R_205_ is a functional important residue that is 20.6L away from the active site S_70_. B. Community 7 of SHV-1, containing 34 residues and is centred by N_136_.

### DyNoPy detects residue couplings essential for protein stability

DyNoPy identifies residue couplings critical for protein functional motions, particularly associated with protein stability. These residue pairs exhibit strong relationships as they are not only directly coupled with each other, but also forms various indirect couplings via other residues. As a result, both residues are considered as core residues inside these communities. It is expected that disruption of these couplings through mutation could compromise collective motions essential for enzyme activity.

As the secondary core residues in community 1 (Fig. 3A) F_72_ is showing a strong coupling with the primary core residue L_162_ and also forms nine indirect couplings with L_162_, including via the catalytic K_234_. This network of direct and indirect relationships reveals the importance of F_72_ and L_162_ coupling in maintaining protein functional motions. Interestingly, previous studies identified a small hydrophobic cavity formed by L_162_ and F_72_, together with L_139_, and L_148_, which is essential for the stability of the active site (50). Notably, DyNoPy successfully recovers the key residues of this local hydrophobic cavity (L_162_, F_72_, and L_148_).

The strong interplay between V_103_ and S_106_, which are both residues on the α3-α4 loop, is seen in community 5 (Fig. 3C). These residues not only interact with each other directly but are also indirectly coupled via 22 other residues. This community emphasizes the significance of hydrophobic nodes in SHV stability and dynamics. Within the analysed 48 residues, 27 are hydrophobic, out of which 15 residues act as nodes critical for enzyme stabilization. Hydrophobic nodes stabilize their own secondary structures and interconnect to stabilize the overall protein (57). V_103_ and S_106_ themselves are hydrophobic nodes, stabilizing α3 helix and α4 helix respectively, and are strongly coupled with each other. In CTX-M, another class A enzyme, N_106_S is a common substitution that results in improved thermodynamic stability and compensate for the loss in stability of the variants (58). Interestingly, this residue is already a Serine in SHV, but still implies its pivotal role in protein stability.

### DyNoPy provides valid explanations for mutation sites

During the evolution of β-lactamases, single mutations on specific sites that are distant from the functional sites have been observed to significantly alter protein catalytic functions. Additionally, single mutations on some surface exposed residues can dramatically increase protein stability. Understanding how these distant mutations impact function and stability becomes a major challenge in understanding protein evolutionary pathways. Communities extracted by DyNoPy show these residues linked with functional important residues, providing a rational for these mutation sites with unknown functions.

Mutations of G_156_ are limited but they lead to ESBL phenotype in the SHV family (46). G_156_ is the central residue for community 4 (Fig. 3B), but it is distant from the active site, over 20L away from the catalytic serine S_70_. Clinical variant SHV-27, has extended resistance ability towards cefotaxime, ceftazidime, and aztreonam (59). It differs from SHV-1 by single amino acid substitution G_156_D, suggesting that it has directly evolved from SHV-1 (59). Limited research has been done on position G_156_, and the understanding of how it affects the enzyme catalytic properties given that it is far away from the active site is still unclear. Based on our results, we suggest that this residue is essential for the overall protein function because of its 11 coevolved dynamic couplings with protein dynamics, including A_146_, another ESBL substitution site.

SHV-38, another ESBL that is capable of hydrolysing carbapenems, harbours a single A_146_V substitution compared to SHV-1 (60). Like G_156_, A_146_ is 16.8 L away from S_70_ but shows an ability in altering protein catalytic function. The A_146_-G_156_ residue pair shows a strong coevolutionary signal and strong correlation with protein overall dynamics, implying that there may compensatory mutations at these sites with potential to emerge in the SHV family in the future. These two residues are not connected to any catalytic residues but their coupling to functional dynamics can offer plausible explanation to ESBL activity of these two mutations.

Unlike other substitution sites that are adjacent to the active site, R_205_ is situated more than 20 L away from catalytic serine S_70_. Its side chain points outward from the protein, exposing to the solvent. The R_205_L substitution often co-occurs with other ESBL mutations and is thought to indirectly contribute to the ESBL phenotype by compensating for stability loss induced by other mutations (61). SHV-3 is an ESBL that exhibits significant resistance to cefotaxime and ceftriaxone (62). Two substitutions in this enzyme, R_205_L and G_238_S, extend its resistance profile (62). Thus, it is promising to see that DyNoPy detected these two mutation sites together within community 6 (Fig. 4A).

Y_105_ and R_266_ are the core residues for community 6. Y_105_ is situated on the α3-α4 loop positioned at the left side of the binding pocket. It is an important catalytic residue that recognizes and binds to the thiazolidine ring of penicillins or β-lactamase inhibitors (63). There is very limited information on the role of R_266_, except that it may stabilize the Ω-loop in the SHV family similar to the analogous T_266_ in TEM (64). G_238_ is coupled with an essential catalytic residue Y_105_, which further links with other catalytic functional residues: S_70_ and A_237_, and R_266_, a residue that known to stabilize the Ω-loop. This indicates that mutations on G_238_ would result in an alteration on protein catalytic function, as well as an increased flexibility of the protein, which strongly aligns with previous finding (62). Its linked mutation site R_205_ does not showing direct coupling with any catalytic residues. Instead, it is directly coupled with R_266,_ which we mentioned as an Ω-loop stabilizer. Thus, it is not surprising that R_205_ substitution alone is never observed in nature (65), as it would not give significant evolutionary advantage to the protein.

### Insights into unexplained functional sites of PDC-3

Unlike the extensively studied SHV-1, the functional roles of individual amino acids in PDC-3 remains largely unexplored. This gap in understanding serves as welcome challenge for interpreting the effects of mutations and the dynamic behaviour of PDC-3 from our results. Although several mutation hotspots, such as those on the Ω-loop (48), have been identified, very little is known about the specific contributions of individual amino acids on the functionality of PDC-3.

In PDC-3, mutations have primarily been reported in the Ω-loop. They enhance its flexibility to accommodate the bulky side chains of antibiotics, while deletions are more common in the R2-loop (43). DyNoPy detected five communities in total (Supplementary Fig. S4B) with all the four predominant Ω-loop mutations appeared in these communities. Community 3, 4 and 5 are explained in the Supplementary information (Fig. S7). Furthermore, DyNoPy also detected several previously unexplored Ω-loop residues.

G_214_, a known mutation site in PDC-3, is the core residue in community 1. Another two essential mutation sites: E_219_ and Y_221_ also participate in this community, directly coupled with G_214_ (Fig. 5A). G_214_ also has direct couplings with four other Ω-loop residues: A_195_, A_197_, G_212_, and L_216_. Previous results have demonstrated that substitutions of Glycine to Alanine or Arginine at 214 significantly destabilizes the Ω-loop (36). The strong correlation between G_214_ and these Ω-loop residues emphasizes the significant contribution of G_214_ towards the stability of the Ω-loop, which corroborates with previous results (36). Moreover, substitutions such as G_214_A and G_214_R and mutations on E_219_ and Y_221_ do not affect R2 loop flexibility, resulting in the smaller active site volume among variants (36) because none of the residues from the R2 loop are detected in this community offering plausible explanation to previously unexplained phenomenon.

**Fig. 5 –.**
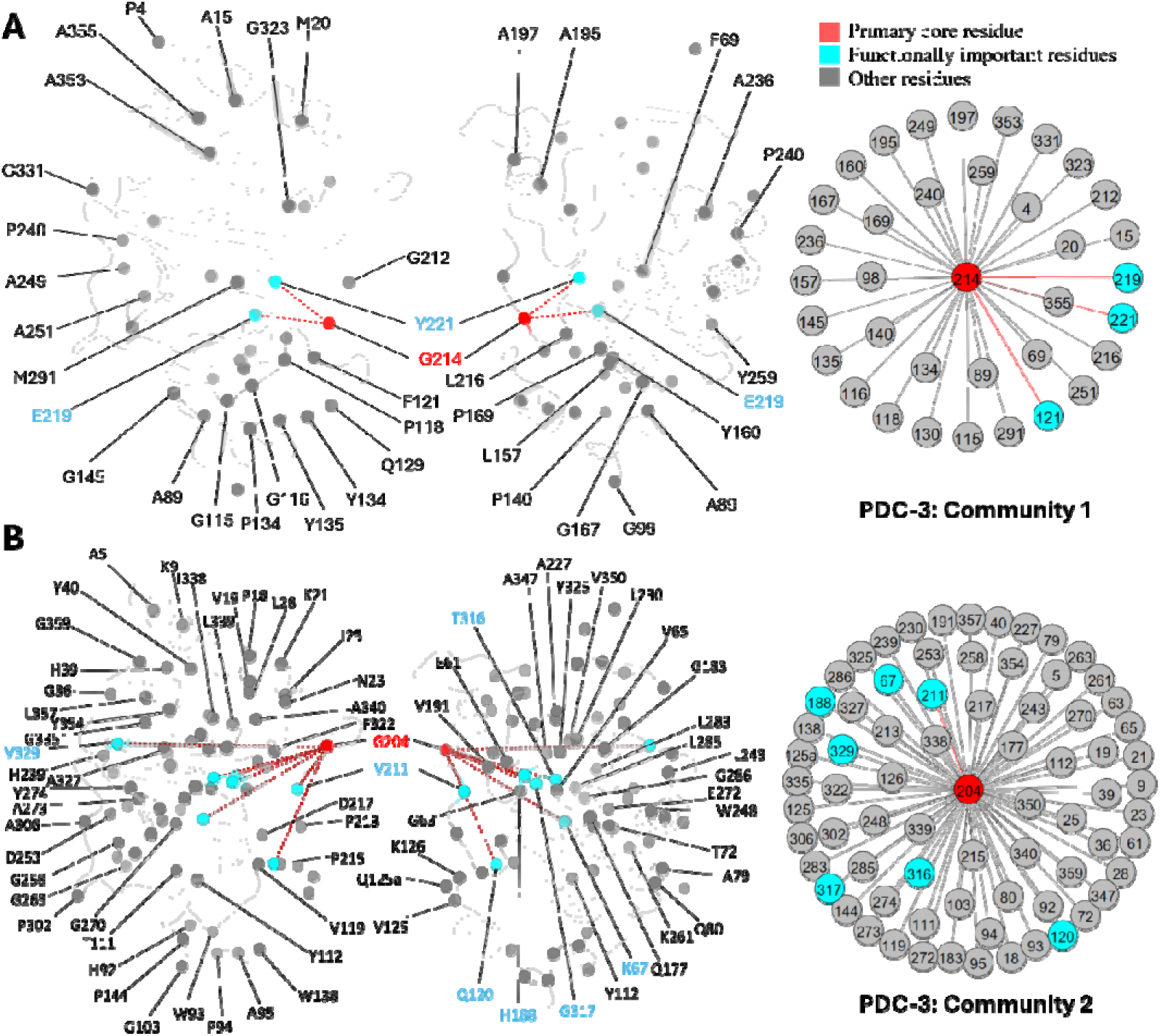
Community 1 and 2 of PDC-3 β-Lactamase. All the residues are depicted as spheres on the protein structure. The core residue for each community is highlighted in red. Functional important residues are marked in cyan. A. Community 1 of PDC-3, comprising 36 residues with G_214_ being the primary core residue. B. Community 2 of PDC-3, containing 74 residues and is centred by G_204_.

G_204_ is the core residue of community 2, coupled with 73 other residues, most of which are distant from the catalytic site, suggesting plausible crucial role in overall protein stability like L_162_ in SHV-1 (Fig. 5B). G_204_, a newly emerged mutation site in the PDC family (66), is located on the short β-sheet β5a within the Ω-loop, near the hinge region between β8 and β9 just above the active site. The only known variant of G_204_ is PDC-466, which was derived from PDC-462 (A_89_V, Q_120_K, V_211_A, N_320_S), with an addition of G_204_D (66). Coupling of G_204_ to several catalytically important residues, including K_67_, K_315_, and T_316_ can suggest that mutations at this site can negatively impact catalytic power. This offers a plausible explanation of seeing fewer variants at this site and mutations at this site could have impact on hydrolysing capabilities of PDC variants. This should be confirmed by further experimental studies of variants of G_204_. Unlike G_214_, E_219_ and Y_221_ mutations which do not influence the dynamics of the R2 loop, substitutions on V_211,_ a member of Ω-loop, has impact on dynamics of R2 loop because of its indirect couplings, through G_204_ to R2-loop residues (36). Two less critical substitution sites, H_188_ and V_329_, were also observed in community 2.

## Conclusions

DyNoPy offers two distinct advantages over existing computational tools (24, 26): a) information on residue-residue coevolution can be directly used to detect the components of protein dynamics that have been preserved during evolution b) dynamic descriptors extracted from the MD ensembles can be used to identify the function-specific conserved dynamic couplings. These couplings are then easily modelled as a graph and network analysis is used to extract epistatic communities and assign roles to residues based on their importance in the graph model. The choice of a relevant descriptor of functional dynamics has an impact on the ability to detect couplings that are involved in functional dynamics. In systems where there is limited role of dynamics in the function, the analysis done with DyNoPy is equivalent to conventional coevolution analysis, which can be consider one limitation of our method.

Here we demonstrated how the choice of relevant global and local descriptors returns a higher number of effective couplings (greater than 0), and in turn leads to interpretable graph models and communities. In other systems, when multiple descriptors can be used to quantify functional conformational change, it is expected that they will differently modulate the effect of coevolution coupling, which will be reflected in a different structure of the associated graph models. This suggests the use of DyNoPy to generate comparative models in proteins with multiple functions associated to distinct dynamical changes.

Mutations of L_162_ and N_136_ have not yet emerged in SHV-1, but they are detected by DyNoPy as core residues for communities. These residues are strongly coupled with other functional important residues, which play critical roles in protein stability and catalytic activity. The identification of these couplings shows high consistency with previous studies and highlights the importance of L_162_ and N_136_ in SHV-1 functional dynamics. Given their central role in these communities, mutations in L_162_ and N_136_ can significantly alter protein function, suggesting their potential for future evolutionary changes. However, their strong relationships with these critical functional residues also suggest that mutation at these sites would need to be balanced to maintain protein function, providing an explanation for why such mutations have not yet emerged in SHV-1 (67). The ability of DyNoPy in detecting functionally important mutation sites was demonstrated via well-characterized mutation sites including R_205_ and G_238_ from SHV-1. Moreover, DyNoPy shows predictive ability on less-studied mutation sites such as G_156_ and A_146,_ by detecting critical residue couplings that coevolved with functional motions.

Based on the knowledge we have gained from analysis of SHV-1 functional protein dynamics we suggest that in PDC-3, mutations at G_204_ because of its significant conserved dynamic couplings can lead to new ESBL/IRBL clinical variants. We suggest that DyNoPy can be used as a predictive tool to identify potential functional residues within this enzyme and guide future mutagenesis studies.

In summary, by integrating hidden evolutionary information with direct dynamic interactions, DyNoPy provides a powerful framework for identifying and analysing functional sites in proteins. The tool not only identifies key residues involved in local and global interactions, but also improves our ability to predict silent residues with previously unknown roles for future experimental testing. Our application of DyNoPy to broad-spectrum β-lactamases ESBLs and IRBLs demonstrates its potential to address key medical challenges such as antibiotic resistance by providing valid predictions on protein evolution.

## Methodology

DyNoPy generates a graph representation of the protein structure that captures the couplings between amino acid residues contributing to the functional dynamics of the protein. Residues are represented as graph nodes, and conserved dynamic couplings are recorded as edges. Edge weights quantify the strength of these couplings. The model is built on two assumptions: residue pairs should have i) coevolved and their ii) time-dependent interactions correlate with a functional conformational change.

Therefore, edge weights (*J_ij_*) for residue *i* and *j* are calculated as:

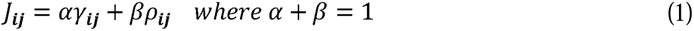

where γ*_ij_* is the scaled coevolution score and *ρ_ij_* is the degree of correlation with the selected functional conformational change. α and β are weights assigned to γ*_ij_* and *ρ_ij_* that have a sum of one. The relative weight of the scaled coevolution score (α) is set to 0.5 in this study. When either of the assumptions listed above is not met, *J_ij_* is set to zero.

### Scaled coevolution scores

The occurrence of residue-residue coevolution can be estimated and quantified using probabilistic models of correlated mutations from deep multiple sequence alignments (MSA). DyNoPy supports generation of the MSA using the HH-Suite package (68) and calculation of scaled coevolution score (γ*_ij_*) using CCMpred (69) as per the protocol described in Bibik et al. (70). For SHV-1 and PDC-3 hhblits returned 18,174 sequences (N_eff_: 11.082) and 27,892 sequences (N_eff_: 9.951). Sequences were detected from the UniRef30 (v2022_02) database (71). First a pairwise residue coevolution matrix (***C***) is calculated, then these raw scores (*C_ij_*) are divided by the matrix mean (Equation 2). All scores (*S_ij_*) smaller than 1 are set to zero, and the remaining values are normalised by the maximum value (Equation 3):

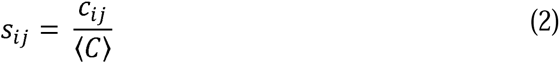

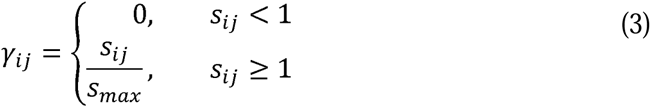

### Correlation with functional motions

The contribution of a residue pair to a selected functional motion is estimated by how much the change in interaction energy between the two residues over time is correlated with a collective variable (CV) describing the functional motion:

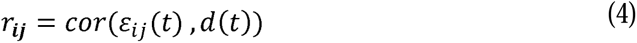

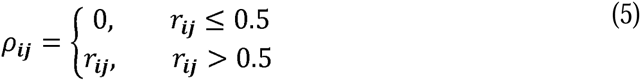

where *ε_ij_*(*t*) is the pairwise non-bonded interaction energy (see details in Supplementary Information) and *d*(*t*) is the time-dependent value of the CV. Examples of CV and a discussion on the choice of the most relevant CV is presented in the results section. Correlation values smaller than 0.5 are set to 0. In absence of detectable contributions to the functional dynamics of the system, the couplings extracted by DyNoPy will describe a pure evolutionary model, and the community detection method presented below will be equivalent to a direct decomposition of the residue coevolution network into units.

### Graph representation and analysis of conserved dynamic couplings

All pairwise conserved dynamic couplings (Equation 1) are collected into a square matrix ***J***. A graph is built from ***J***, using python-igraph v0.11 library (72). Nodes represent residues, and edges are drawn between nodes with positive *J_ij_*. Edge weights are set to *J_ij_*. The relative importance of the residues in this model of protein dynamics is calculated as eigenvector centrality of the nodes (73). Residues involved in extensive correlated dynamics with other highly connected residues have higher eigenvector centrality (EVC) scores. Groups of residues contributing to important collective motions are detected by community analysis of the graph structure. The Girvan-Newman algorithm is used to extract the community structure (74). A meaningful community should contain at least three residues. Applying network analysis on the combined dynamics-coevolution matrix helps us extracting higher-order interactions beyond pairwise coupling and detecting critical residues, which show multiple interactions with each other. Moreover, indirect long-range relationships, which would be hard to identify from numerical data, could be detected through community clustering. Community-based analysis offers a more comprehensive understanding of residue relationships and enables the visualization of residue couplings on the protein structure.

### Adaptive Sampling Molecular Dynamics Simulations

MD simulation data was sourced from our previous studies (35, 36). To summarise, SHV-1 structural coordinates (PDB ID: 3N4I) were obtained from the Protein Data Bank and modified to the wild type by introducing the E104D mutation. Similarly, the PDC-3 structure was derived from PDC-1 (PDB ID: 4HEF) by a T105A substitution. Both enzymes were protonated at pH 7.0 using PropKa from the PlayMolecule platform (75). One disulfide bond between C_77_ and C_123_ was specified in SHV-1. Both structures were solvated with TIP3P water molecules in a periodic box with a box size of 10 Å. Ions were added to neutralize the overall charge of each system at 150mM KCl. Amber force field ff14SB was used for all MD simulations (76). After an initial minimisation of 1000 steps, both the enzymes were equilibrated for 5 ns in the NPT ensemble at 1 atmospheric pressure using the Berendsen barostat (77). The initial velocities for each simulation were sampled from the Boltzmann distribution at 300 K. Multiple Markov State Model (MSM)-based adaptively sampled simulations were performed for both proteins based on the ACEMD engine (78, 79). A canonical (NVT) ensemble with a Langevin thermostat (80) (damping coefficient of 0.1 ps−1) and a hydrogen mass repartitioning scheme were employed to achieve time steps of 4 fs. For SHV-1, each trajectory spanned 60 ns with a time step of 0.1 ns, with a total of 593 trajectories. In the case of PDC-3, 100 trajectories were collected, each containing 3000 frames, lasting 300 ns. To manage the extensive datasets efficiently, trajectories were strategically stridden to ensure that a minimum of 30,000 frames were preserved for each system. The resulting trajectories are summarized in Supplementary Table S4.

### Calculation and Selection of Collective Variables

DyNoPy works on the assumption that time-dependent interactions between critical residues, either having significant structural change or not will correlate with functional conformational motions. Since MD simulation data is high-dimensional, a time-dependent collected variable (CV) is required to extract the most relevant information for the process under study. The usefulness of DyNoPy is dependent on the choice of the CVs. To guide the selection of CVs, we selected 12 distinct features: radius of gyration (R_g_), the first principal component (PC1), partial PC1 (PC1_partial), the first time-lagged independent component (TC1), partial TC1 (TC1_partial), global root mean square deviation (gRMSD), partial RMSD (pRMSD), dynamical RMSD (dRMSD), global solvent accessible surface area (gSASA), partial SASA (pSASA), active site pocket volume, and the number of hydrogen bonds (hbond). A description of the CVs, including the calculation methods and the residues used to calculate the partial variables, is detailed in the Supplementary information. CVs were subsequently used as input features for DyNoPy. A good collective variable (CV) should appropriately describe protein functional motions. Thus, a CV that detects the highest number of residue couplings is expected to be the most suitable descriptor. The length of the MD simulations should be appropriate to effectively sample the desired functional process as described by the selected CV.

## Data Availability

All files required to run the simulations (topology, coordinates, input), processed trajectories (xtc), corresponding coordinates (pdb), can be downloaded from the DOI https://doi.org/10.57760/sciencedb.15876 (PDC-3) and 10.5281/zenodo.13693144 (SHV-1). DyNoPy is available at https://github.com/alepandini/DyNoPy.

## Acknowledgements

SCD was supported by Leverhulme Trust grant RPG-2017-222 awarded to AP and JAG. The authors would like to thank Arianna Fornili for insightful suggestions on the design of DyNoPy methodology.

