## Supplementary Information for "Functionally Important Residues from Graph Analysis of Coevolved Dynamic couplings"

***Corresponding author:**

Shozeb Haider

ORCID: 0000-0003-2650-2925

Alessandro Pandini

ORCID: 0000-0002-4158-233X

**This PDD file includes:**

Supporting text
Figures S1 to S7
Table S1 to S4
SI References

### Supporting Information Text

#### Interaction energies (E)

The pairwise non-bonded interaction energies for upper triangle of the NxN residue matrix is calculated using dyno_pwie.py which is a wrapper to ccptraj from ambertools. The list of pairs is generated and depending on the number of available threads ($n_t$), '$n$' sets of pairs are created, and each pair set is assigned to an instance of ccptraj for calculation of pairwise interaction energies. As this is a RAM dependent operation, based on the size of the MD trajectory, care must be taken to not spawn too many instances of ccptraj as it will slow down each instance of the ccptraj calculation. For each residue pair ($r_{ij}$), a separate file with coulomb ($q_{ij}$) and Van Der Waals ($w_{ij}$) interaction energies, as two separate columns, are saved to a h5py compressed file and the total interaction energy ($\epsilon_{ij}$) for each time step can be calculated from this data using equation (1).

$$\epsilon_{ij} = q_{ij} + w_{ij} \quad (1)$$

#### Collective Variables (CVs) Description

#### Radius of Gyration (Rg)

The radius of gyration (Rg) measures the compactness of the protein structure through the calculation of the root mean square distance of all atoms from the centre of mass. A lower Rg indicates a more compact structure, while a higher Rg suggests a more expanded or unfolded state. This CV was calculated using the MDAnalysis v2.7.0 (9, 10), focusing on the whole protein for both enzymes.

#### The first Principal Component (PC1)

Principal Component Analysis (PCA) identifies the major dynamical feature of a protein by reducing the dimensionality directly from molecular dynamics simulation data. PC1 represents the largest variance on selected features across the simulation.

Global PC1 was computed using PyEMMA v2.5.12 (11), focusing on backbone torsion angles and $\chi 1$ angles for all residues, capturing the global dominant structural changes.

**The partial PC1 (PC1_partial)**

PC1_partial was calculated using the same methodology as PC1 but only focusing on functionally significant residues and essential secondary structures. This approach prioritizes the dynamics within regions critical to the protein's function, thereby providing a more targeted analysis of pertinent conformational changes. It is particularly well-suited for studying proteins that are predominantly rigid with localized flexible regions.

**Time-lagged Independent Component 1 (TC1)**

Time-lagged Independent Component Analysis (TICA) identifies slow, independent processes within the protein dynamics. TC1 captures the slowest conformational changes in the protein over time. It was computed globally using backbone torsion angles and $\chi 1$ angles for all residues, with PyEMMA v2.5.12 (11).

**Partial Time-lagged Independent Component 1 (TC1_partial)**

Similarly, TC1_partial was calculated, but focused on specific, functionally important regions. By analysing these key residues and structures, TC1_partial represents the slow conformational changes crucial within essential regions, which are highly corresponding with protein functions.

**Global Root Mean Square Deviation (gRMSD)**

The RMSD measures the average deviation of a protein's atomic positions from a reference structure, typically the starting conformation. gRMSD provides an overview of the protein's structural deviation over time. In our case, trajectories were first aligned, and Cα RMSD was calculated using MDAnalysis v2.7.0 (9, 10), utilizing the starting conformation of each protein as a reference.

**Partial Root Mean Square Deviation (pRMSD)**

This CV is calculated similarly to gRMSD but focuses on specific residues or regions of interest. pRMSD provides insight into the structural stability of functionally important areas of the protein by focusing on catalytical residues and essential loops.

**Dynamical Root Mean Square Deviation (dRMSD)**

dRMSD is an extension of RMSD that considers the dynamic nature of protein motions by analysing fluctuations over time rather than static deviations. More specifically, instead of calculating RMSD to a fixed reference structure, it focuses on the difference between adjacent frames. This approach allows the analysis of short-timescale fluctuations and provides insight into the dynamic nature of protein motions over time. It was also calculated using MDAnalysis v2.7.0 (9, 10) focusing on the Cα and offers a more nuanced understanding of the protein's conformational landscape.

**Global Solvent Accessible Surface Area (gSASA)**

SASA quantifies the surface area of the protein accessible to the solvent. gSASA was calculated using GROMACS v2020.1 (12) and provides information about the protein's overall exposure to the solvent, which is relevant for understanding folding and binding interactions.

**Partial Solvent Accessible Surface Area (pSASA)**

pSASA focuses on the solvent exposure of specific active site residues, offering insights into how binding sites or functional regions of the protein interact with the solvent.

**Active Site Pocket Volume**

The volume of the active site pocket was calculated using Mdpocket (13). This CV is important for understanding the size and shape changes in the binding site, which can influence ligand binding and protein function. Upon importing the topology file and trajectories, Mdpocket autonomously identifies potential binding pockets within the protein structure. Subsequently, the relevant pocket was manually selected (Fig. S3)

**Number of Hydrogen Bonds (hbond)**

The number of hydrogen bonds was calculated per frame using VMD v1.9.3 (14), with a distance threshold of 3.5Å and an angle cutoff of 40°. This CV provides insight into the stability of the protein structure and interactions that are critical for maintaining its conformation and function.

**Residues and Regions Used in Partial CVs**

For SHV-1, the backbone torsion angles and χ1 angles for catalytic important residues: S70, T71, K73, S130, N132, K234, and T235, and Ω-loop residues (R164-D179) were used as the input feature to calculate the partial CVs. Similarly, for PDC-3, the backbone torsion angles and χ1 angles for catalytic important residues: K67, Y150, N152, K315, T316, and G317, Ω-loop residues (G183-S226), and R2-loop residues (L280-Q310) were utilized to calculate all the partial variables.

**Other SHV-1 Communities**

Core residues for some of the communities identified by DyNoPy have never been studied in class A β-lactamases before. DyNoPy predicts these residues as essential due to their coevolution trends with many other residues and their critical role in class A β-lactamase dynamics. The graph depicting the community structure is available in the supplementary materials (Fig. S6).

$D_{267}$ is the core residue in community 3 (Fig. S6A). This residue is located on the loop connecting β-sheet β9 and α-helix α12. It is outside the catalytic site and has not undergone essential substitutions, thus remaining unexplored in class A β-lactamases. However, DyNoPy indicates its potential importance due to its relationships with three known essential mutation sites: $L_{35}$, $E_{240}$, and $A_{187}$. $L_{35}$ and $E_{240}$ are predominant ESBL mutation sites, while $A_{187}$ substitution confers an inhibitor-resistant phenotype (15). The relationship between $D_{267}$ and these mutation sites suggests a trend towards coevolution, and interactions between $D_{267}$ and these mutation sites are important for protein dynamics.

Similarly, $I_{279}$, located also on α12, is the core residue of community 8 (Fig. S6B). This residue forms relationships with $R_{244}$, a critical inhibitor resistant mutation site (16). Other residues within this community are predominantly positioned on the α1 and α12 helices, near the protein's terminus, highlighting the significant contribution of $I_{279}$ to protein integrity and stability.

For Community 9, $I_{155}$ serves as the core residue (Fig. S6C). This community is relatively localized and does not encompass any known essential residues. Most of the residues involved in this community are spatially close to each other, suggesting that $I_{155}$ plays a vital role in the local dynamics, especially surrounding α-helices α7 and α9.

**Other PDC-3 Communities**

The core residues in the other PDC-3 communities have not been extensively studied. However, their detection by DyNoPy suggests a significant trend in co-evolution and highlights their crucial role in protein dynamics (Fig. S7). This emphasizes the capability of DyNoPy to predict essential residues in previously unexplored proteins, potentially offering valuable insights for future experimental research.

$E_{49}$, $D_{206}$ and $R_{210}$ are core residues for community 3, a small community containing only 14 residues (Fig. S7A). $R_{210}$ is the primary core residue in this community, linking to six residues, while both $E_{49}$ and $R_{210}$ are secondary core residues that show a relationship with four residues. Unlike other communities with widespread interactions, community 3 illustrates a localized relationship among Ω-loop residues, primarily on the short β5a and β5b β-sheets and the short helix α7a. 8 out of 14 residues are Ω-loop residues, with the remainder located on adjacent loops or near the terminals of adjoining secondary structures, all of which are flexible regions. This community indicates that $R_{210}$ is crucial for maintaining local structural integrity and the stability of the Ω-loop.

$G_{202}$, the core residue of community 4, appears essential for active site stability by interacting with residues whose side chains point into the active site (Fig. S7B). It also stabilizes two α-helices, α2 and α5, located adjacent to the active site. $P_{154}$ is a special mutation site in the PDC family. $P_{154}L$ occur in PDC-73 and PDC-81, giving the protein a mild increase in resistance to ceftazidime (17).

$K_{204a}$ and $R_{207}$ are the central residues in Community 5, each establishing six interactions and thus sharing an equal position of importance within this community. $K_{281}$ and $K_{351}$, which interact with both core residues, are highlighted in light yellow (Fig. S7C). $K_{204a}$ forms direct interactions with six residues, primarily located on the opposite side of the active site, either on the R2 loop or at the beginning of helix α11. This suggests that $K_{204a}$ plays a crucial role in maintaining the conformation of the R2 site. Additionally, $K_{204a}$ shows direct correlations with the catalytically significant residue $K_{315}$. Although $R_{207}$ is spatially close to $K_{204a}$, it interacts with residues on loops that are distant from the catalytic site.

### Detailed Structure Information for SHV-1 and PDC-3

In class A β-lactamases, the active site is surrounded by three loops: the α3-α4 loop (residues 101-111), the Ω-loop (residues 164-179), and the hinge region (residues 213-218) (3). The Ω-loop is particularly critical as it positions $N_{170}$ to hydrogen bond with the $E_{166}$ via a conserved water molecule, which is essential for initiating the deacylation step (4). Compared to class A β-lactamases, the active site of class C β-lactamases is wider, conferring a broader substrate binding capability (5) (Fig. S3). The active site of class C β-lactamases can be divided into two parts: the R1 site and the R2 site (6). The R1 region is surrounded by the extended Ω-loop (residues 183-226), while the R2 site is enclosed by the R2-loop (residues 280-310) (7). The Ω-loop in class C β-lactamases is significantly longer than that in class A, enhancing the active site ability to accommodate diverse substrates and contributing to the extended spectrum profile of some class C enzymes (7).

**Supplementary Figures**

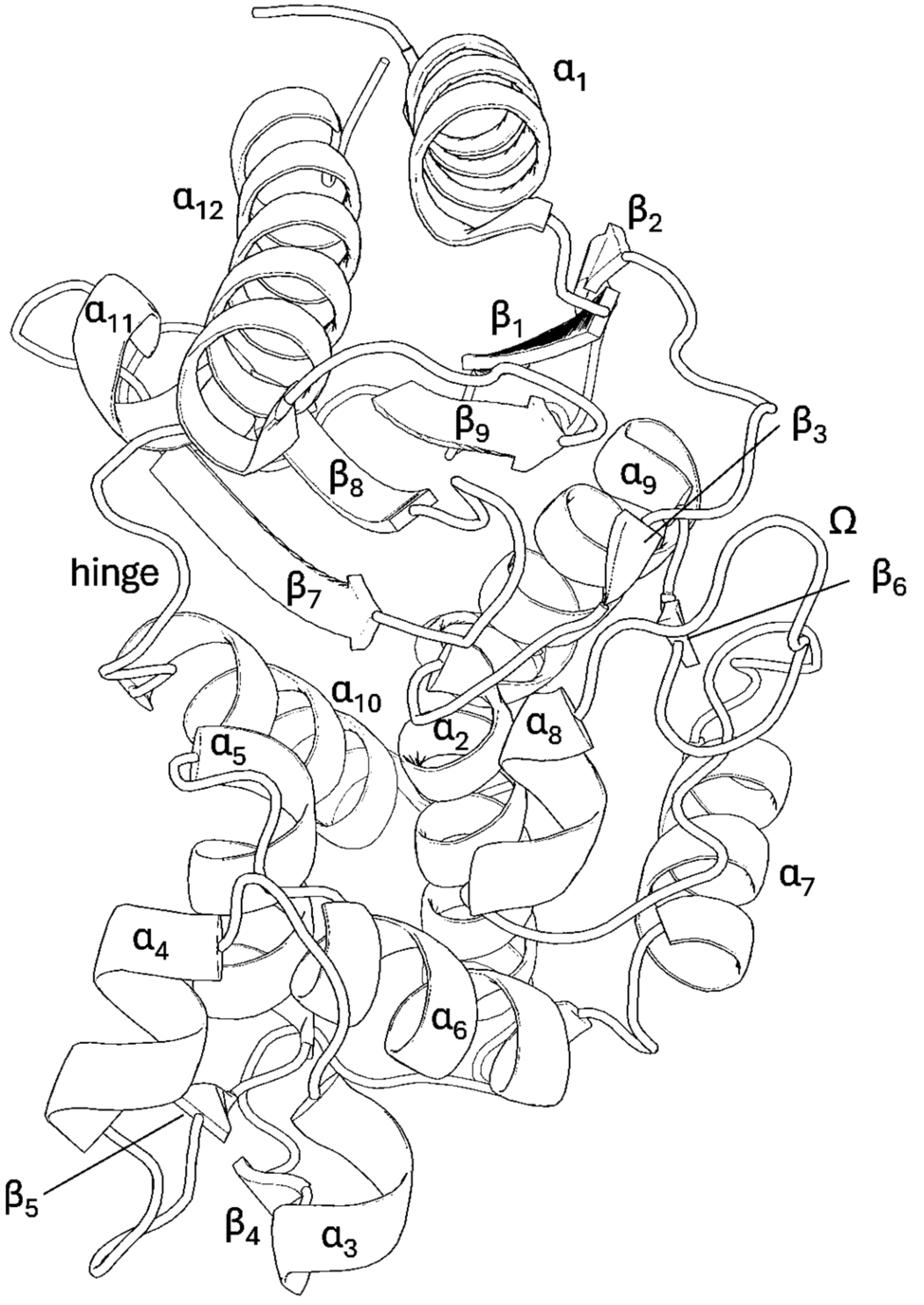


**Fig. S1 | SHV-1 structural nomenclature.** PDB id: 3N4I (1).

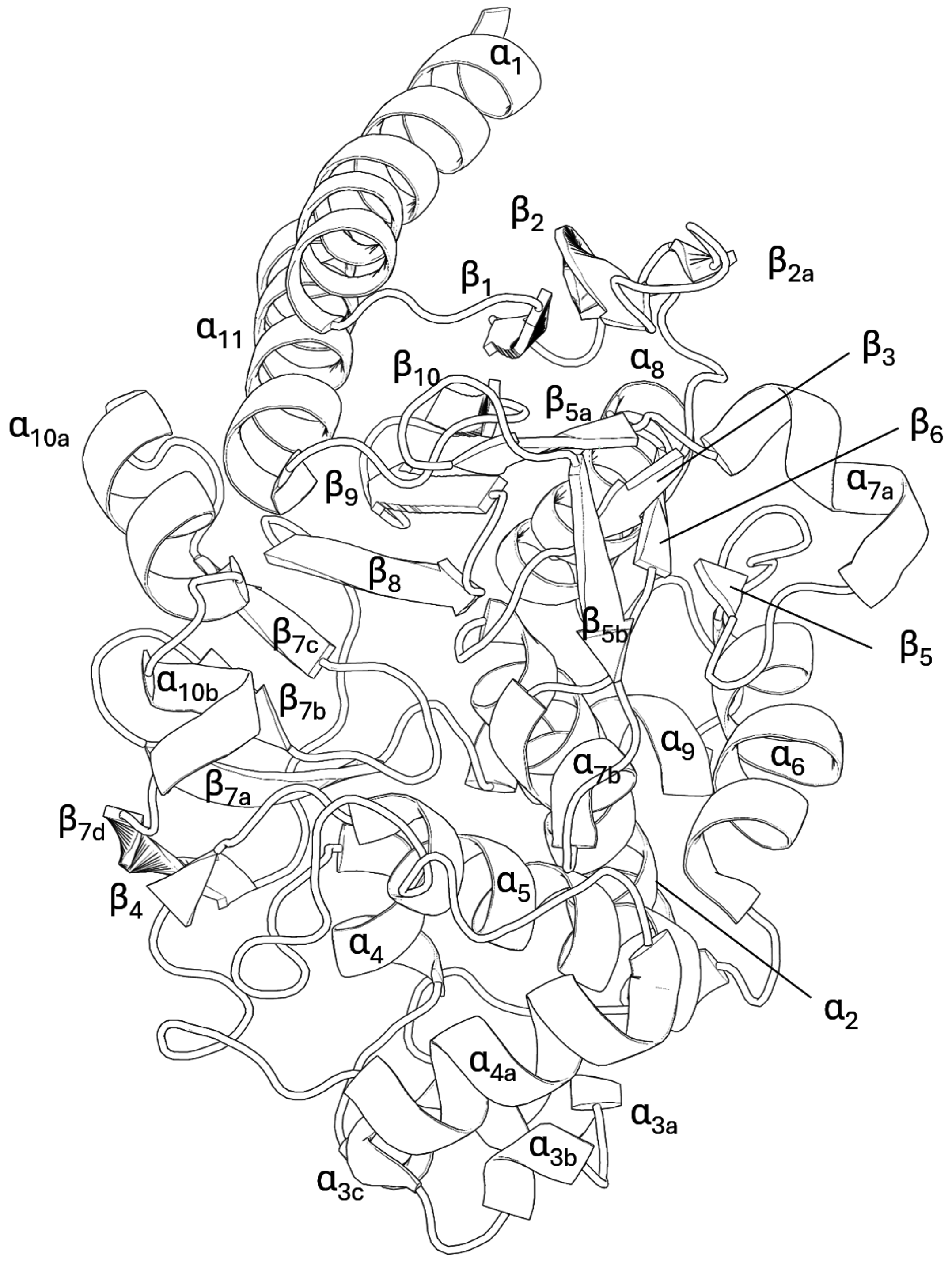


**Fig. S2 | PDC-3 structural nomenclature.** PDB id: 4HEF (2).

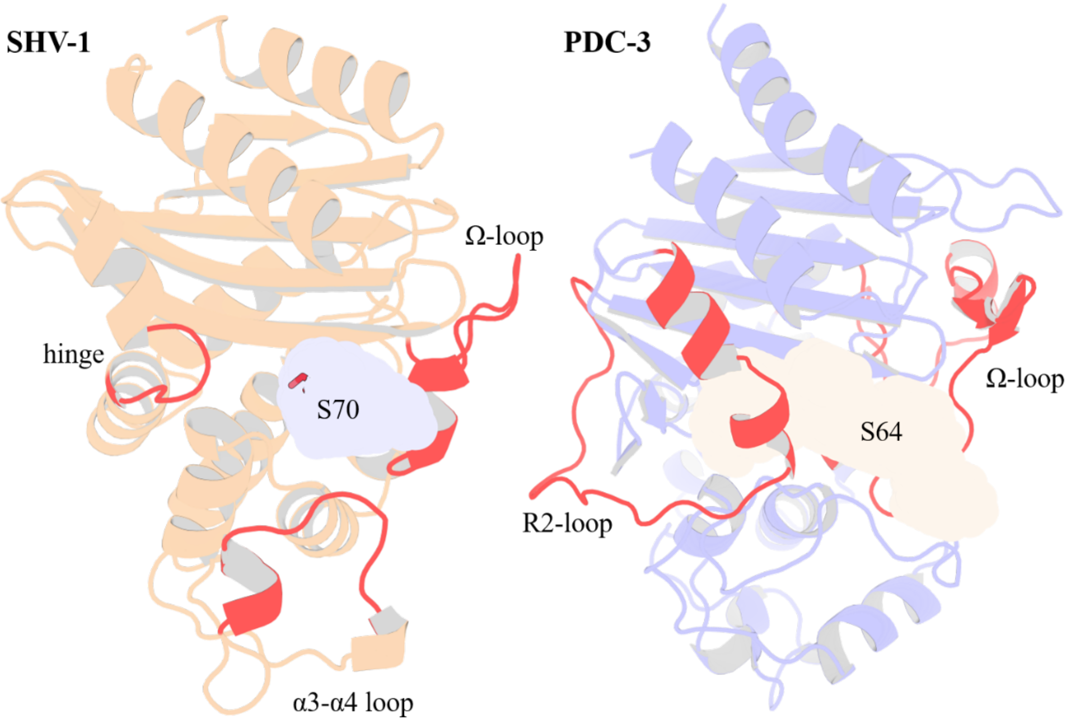


**Fig. S3 | Crystal structure and binding pocket of SHV-1 (PDB id: 3N4I) and PDC-3 (PDB id: 4HEF).** $S_{70}$ and $S_{64}$ are the catalytic serine residue for SHV-1 and PDC-3 respectively. The α3-α4 loop, the Ω-loop, and the hinge region in SHV-1 are highlighted in red. The Ω-loop and the R2-loop in PDC-3 are colored in red.

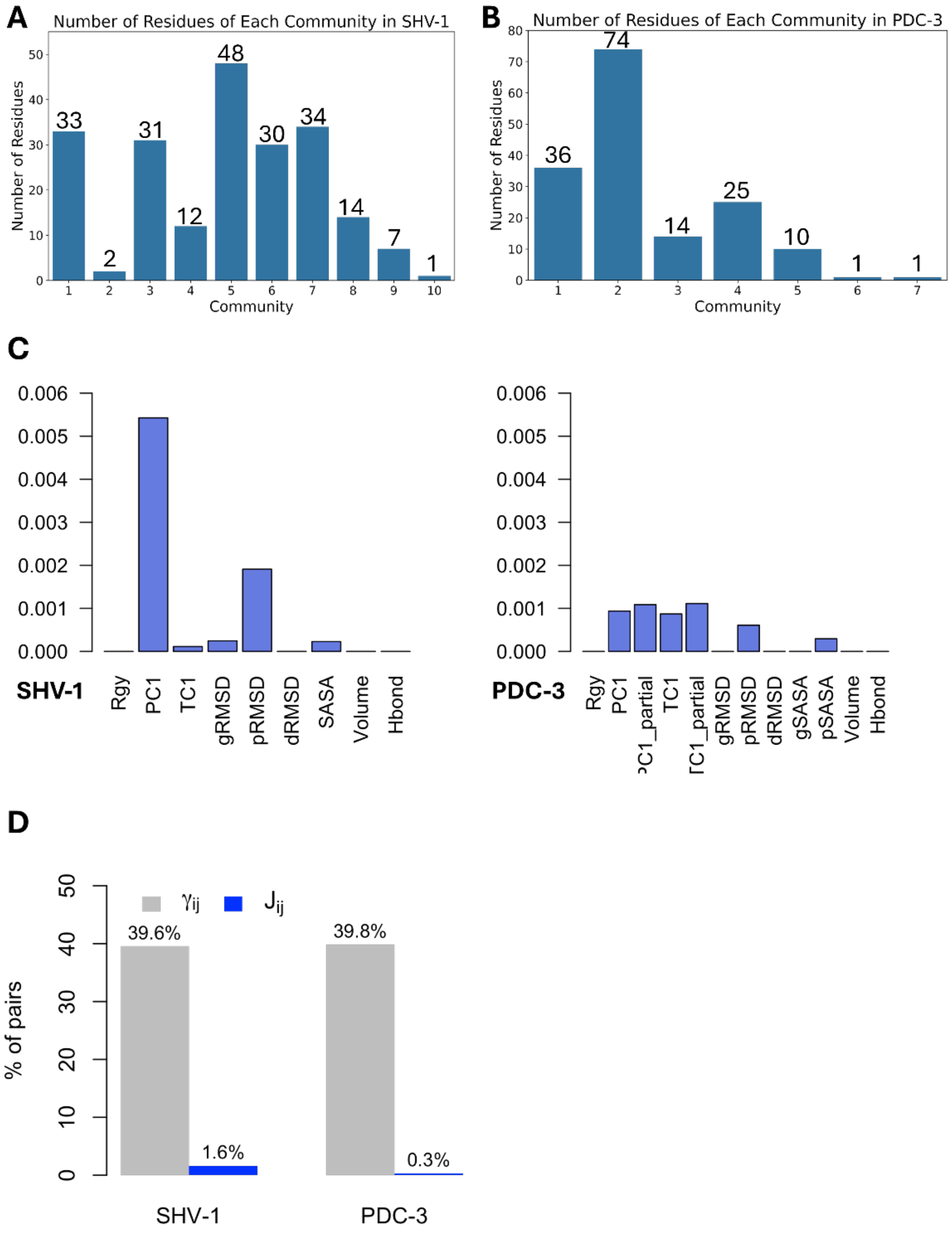


**Fig. S4** | **A.** Number of residues of each community in SHV-1. **B.** Number of residues of each community in PDC-3. A reasonable residue community should contain at least three residues. **C.** Average *J* matrix score varies across different CVs. Left: SHV-1; Right: PDC-3. **D**. Number of non-zero couplings detected by scaled coevolution scores ($\gamma_{ij}$) and *J* values calculated by DyNoPy ($J_{ij}$).

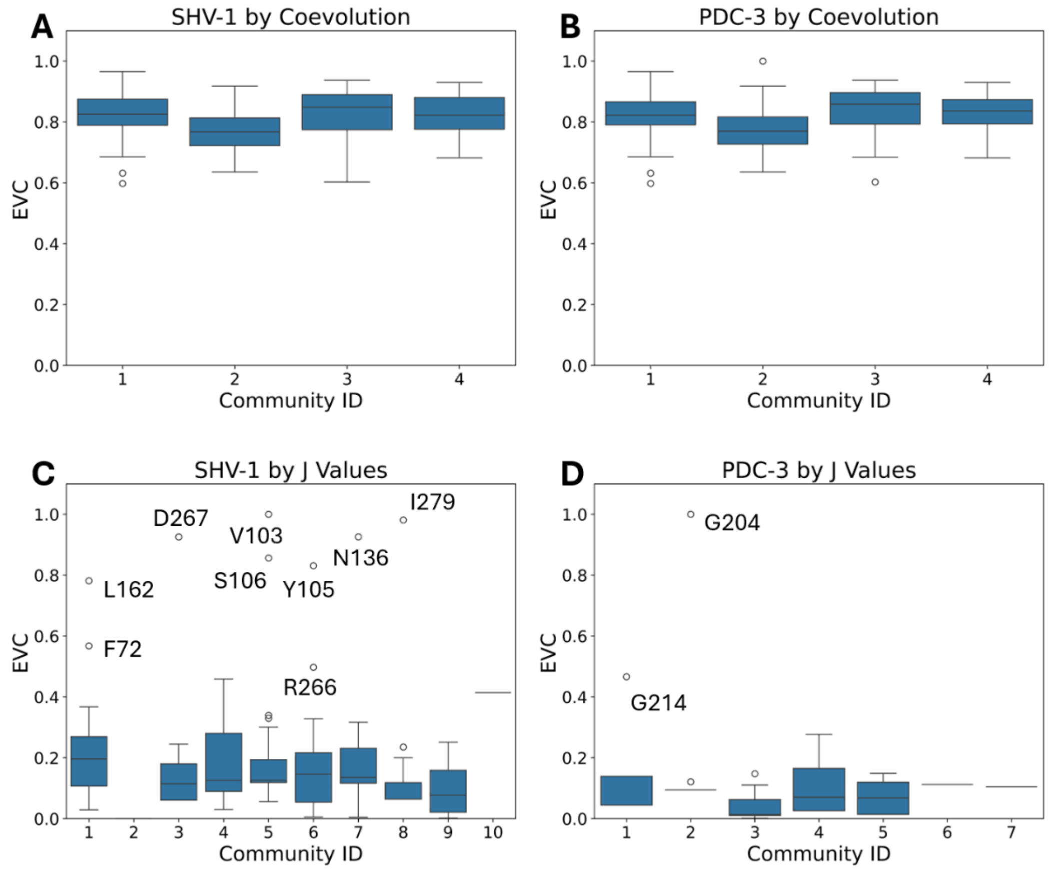


**Fig. S5 | Eigenvector Centrality (EVC) distribution of all the residues for each community. A.** EVC distribution of SHV-1 for each community detected by traditional coevolution scores. **B.** EVC distribution of PDC-3 of each community detected by traditional coevolution scores. **C.** EVC distribution of SHV-1 of each community induced from J value calculated by DyNoPy. **D.** EVC distribution of PDC-3 of each community induced from *J* value calculated by DyNoPy.

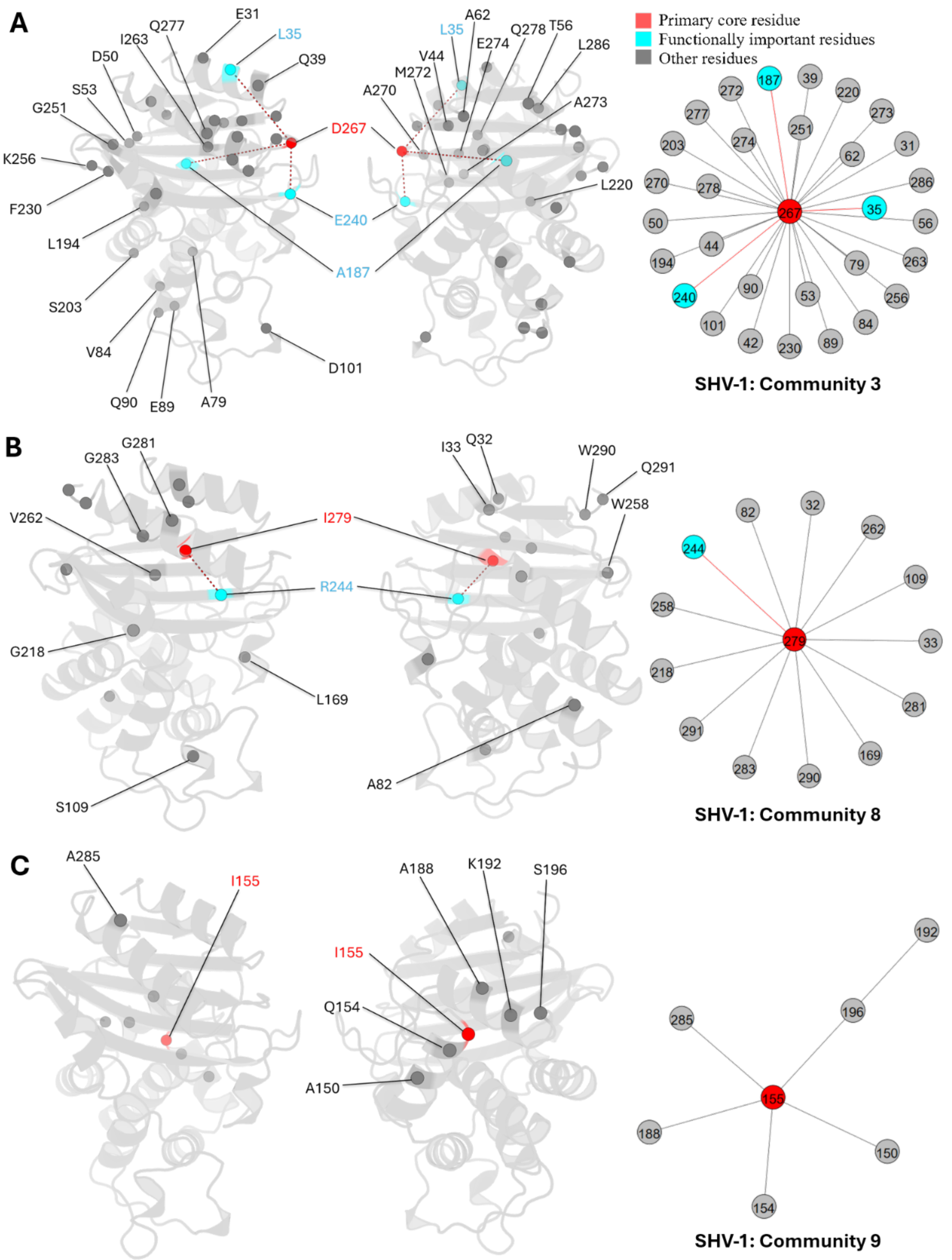


**Fig. S6 | Community 3, 8 and 9 of SHV-1 β-Lactamase.** All the residues are depicted as spheres on the protein structure. The core residue for each community is D267, I279, and I155, respectively. They are highlighted in red. Functional important residues are marked in cyan. A. Community 3 of SHV-1, comprising 31 residues. B. Community 8 of SHV-1, containing 14 residues. C. Community 9 of SHV-1, with 7 residues.

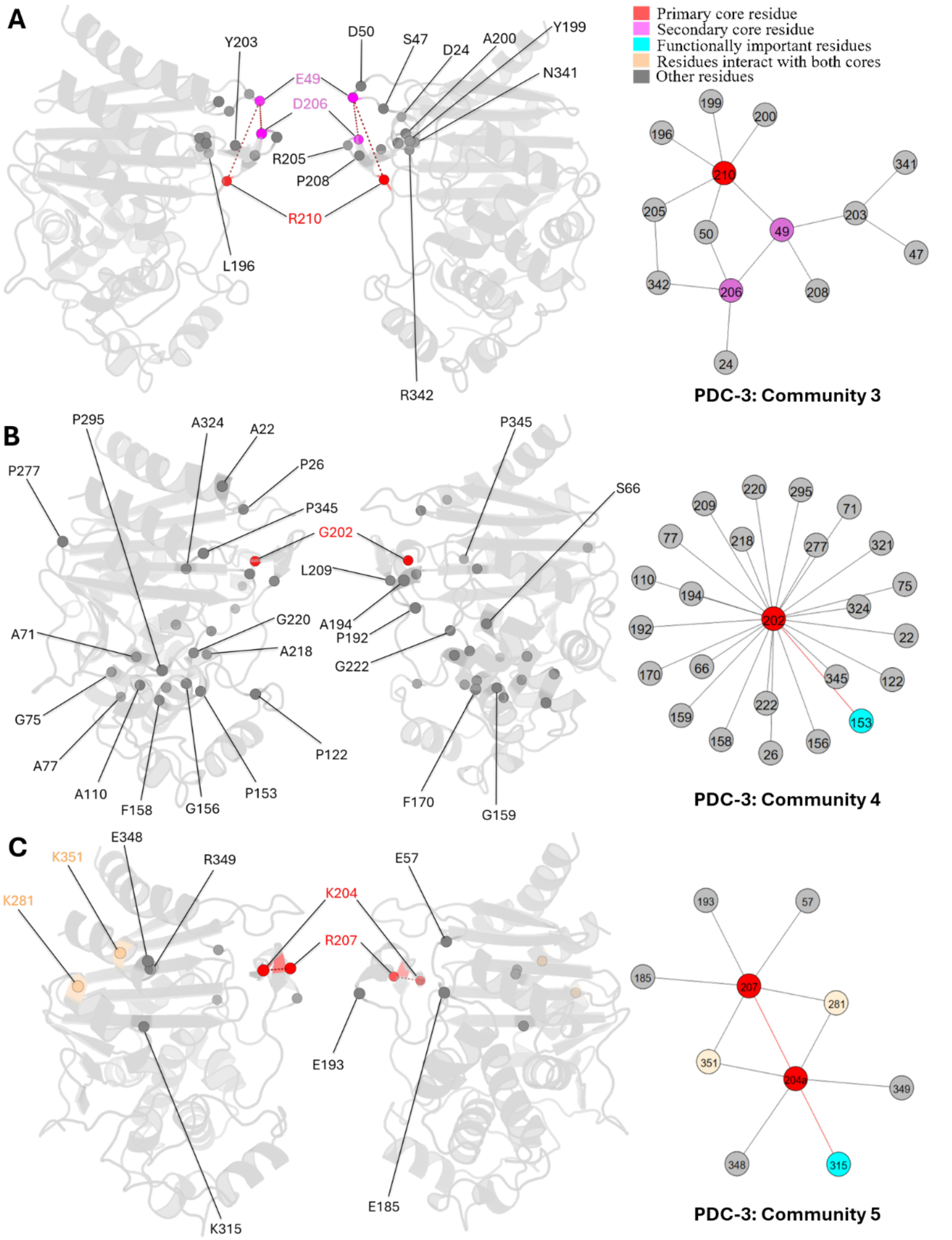


**Fig. S7 | Community 3, 4 and 5 of PDC-3 β-Lactamase.** All the residues are depicted as spheres on the protein structure. The core residue for each community is highlighted in red, while purple is used to emphasize the secondary core residue. Residues that interact with both cores are coloured in light yellow. Functional important residues are marked in cyan. A. Community 3 of PDC-3, comprising 14 residues with $R_{210}$ being the primary core residue. B. Community 4 of PDC-3, containing 25 residues and is centred by $G_{202}$. C. Community 5 of PDC-3, embracing 10 residues and having two core residues $K_{204}$ and $R_{207}$.

**Supplementary Tables**

**Table S1:** Hydrophobic nodes in SHV-1 (8)

| ***Nodes*** | ***Residues*** | ***Residue No.*** |
|---|---|---|
| *α2* | VVLCGAVLA | 74-82 |
| *α3* | DLV | 101-103 |
| *α4* | SPV | 106-108 |
| *α5* | AAAI | 124-127 |
| *α6* | SAA | 133-135 |
| *α6* | LLL | 137-139 |
| *α9* | SMA | 185-187 |
| *α10* | RLSA | 198-201 |
| *α11* | SVL | 223-225 |
| *β7* | FIA | 230-232 |
| *β8* | ALL | 248-250 |
| *β9* | AVV | 260-262 |
| *α12* | AGIG | 280-283 |

**Table S2:** Dynamic descriptors and number of residue pairs detected

| ***Dynamic Descriptors*** | ***Number of Pairs Detected*** | |
|---|---|---|
| | **SHV-1** | **PDC-3** |
| $R_g$ | 0 | 0 |
| *PC1* | 571 | 185 |
| *PC1_partial* | 161 | 211 |
| *TC1* | 119 | 172 |
| *TC1_partial* | 339 | 216 |
| *gRMSD* | 26 | 0 |
| *pRMSD* | 203 | 117 |
| *dRMSD* | 0 | 0 |
| *gSASA* | 23 | 0 |
| *pSASA* | 18 | 62 |
| *volume* | 0 | 0 |
| *hbond* | 0 | 0 |

**Table S3:** Number of residues in each community detected using only the coevolution scores

| | ***Community 1*** | ***Community 2*** | ***Community 3*** | ***Community 4*** |
|---|---|---|---|---|
| *SHV-1* | 84 | 50 | 44 | 86 |
| *PDC-3* | 93 | 57 | 82 | 126 |

**Table S4:** Summary of molecular dynamics simulation systems

| ***System*** | ***Number of Trajectories*** | ***Frames for Each Trajectory*** | ***Total Simulation Time (μs)*** | ***Stride*** | ***Total Frames After Stride*** |
|---|---|---|---|---|---|
| *SHV-1* | 593 | 600 | 35.58 | 10 | 35580 |
| *PDC-3* | 100 | 3000 | 30 | 10 | 30000 |

*All trajectories were simulated with a time step of 0.1 ns
